# Differential gene expression analysis identifies a group of defensin like peptides from *Solanum chacoense* ovules with *in vitro* pollen tube attraction activity

**DOI:** 10.1101/2024.10.25.620243

**Authors:** Yang Liu, Valentin Joly, Mohamed Sabar, Daniel Philippe Matton, David Morse

**Affiliations:** Institut de Recherche en Biologie Végétale, Département de Sciences Biologiques, Université de Montréal, Montréal, Québec, Canada

**Keywords:** attractants, cysteine rich peptides, defensin-like peptides, embryo sac-dependent genes, embryo sac maturation, pollen tube guidance, semi-*in vivo* assays, *Solanum* species

## Abstract

*Solanum chacoense* is a wild potato species with superior genetic resistance to diseases and pests that has been extensively used for introgression into cultivated potato. One determinant of crossing success between wild and cultivated potato species is the effective ploidy of the parents. However, little is known about whether other, prezygotic level, breeding barriers exist. We hypothesize ovular pollen tube guidance may serve as such a checkpoint. Tests for species-specific pollen tube guidance using semi-*in vivo* assays suggested a positive correlation between species-specificity and taxonomic distance. RNA-seq of ovules hand dissected from wild type plants at anthesis and two days before anthesis, as well as from a *frk1* mutant lacking an embryo sac identified a list of 284 embryo sac-dependent genes highly expressed in mature ovules and poorly expressed in all other samples. Among these are 17 *<u>S</u>olanum <u>c</u>hacoense* <u>c</u>ysteine-<u>r</u>ich <u>p</u>roteins (ScCRPs), considered to be good candidates since CRPs are ovular pollen tube attractants in other species. A group of three cloned and purified ScCRP2 sequences belonging to the DEFL protein family showed moderate levels of *in vitro* pollen tube attraction activity in functional assays. We conclude that ScCRP2s are good candidates for ovular pollen tube guidance in *S. chacoense*.

**Highlights:** This work introduces key characteristics of pollen tube guidance in *Solanum chacoense*, identifies candidate attractants through transcriptome subtraction and reports *in vitro* pollen tube attraction activity for three defensin-like peptides.

## Introduction

Pollen tube guidance is a process whereby the direction of pollen tube growth is influenced by molecules secreted from pistil tissues. Based on where the pollen tubes are in the pistil, pollen tube guidance is divided into two phases: pre-ovular guidance (on the stigma and in the style) and ovular guidance (approaching the ovule) (Higashiyama and Takeuchi, 2015). In pre-ovular guidance, some important players include Chemocyanin and TTS, which were identified from lily stigma and tobacco style, respectively (Cheung *et al*., 1995; Kim *et al*., 2003; Wu *et al*., 1995). Both proteins induced positive pollen tube chemotropism (pollen tubes grow towards these proteins) in an *in vitro* assay.

When pollen tubes arrive at the end of the style, they come under the influence of the ovule. Ovular pollen tube guidance has been described as occurring in two distinct steps. First, a long-range guidance was suggested in *Arabidopsis* to describe the ovule-derived activity that attracts pollen tubes to exit the stylar transmitting tract and to approach the septum (Hulskamp *et al*., 1995). Following this long-range activity, pollen tubes transition from an intercellular growth mode (in the transmitting tract) to a surface growth mode, but the molecules involved remain unknown. Second, a short-range guidance was described in which pollen tubes, growing within 100 µm of an ovule, make sharp turns to enter the micropyle of an ovule (Palanivelu and Preuss, 2006). Two discrete phases of short-range guidance were proposed: 1) funicular guidance, that leads pollen tubes to the funiculus, and 2) micropylar guidance, where the pollen tubes reorient to target the micropyle for double fertilization (Shimizu and Okada, 2000).

Some flowering plants have a structure termed obturator at the funiculus which is involved in funicular guidance. An obturator is a protuberance from the funiculus that facilitates pollen tube access to the micropyle. This structure is of particular importance in plants bearing anatropous ovules (where the micropyle is adjacent to the funiculus-placenta junction), as pollen tubes need to be reoriented in order to arrive at the micropyle. For example, in the apple ovary, it was observed that pollen tubes make a sharp 180° turn on the surface of the obturator before entering the micropyle. A direct involvement of obturator secretion in ovular pollen tube guidance has not been addressed, although immunolocalization studies show the obturator in the apple flower secretes arabinogalactan proteins, some of which are developmentally regulated while others are elicited by the presence of pollen tubes (Losada and Herrero, 2017).

Micropylar attractants have been identified in three species. All are small secreted peptides that induce pollen tube chemotropism. They include ZmEA1 in maize, LURE peptides in *Torenia* and *Arabidopsis*, as well as XIUQIU peptides in *Arabidopsis* (Kanaoka *et al*., 2011; Marton *et al*., 2012; Okuda *et al*., 2009; Takeuchi and Higashiyama, 2012; Zhong *et al*., 2019). LUREs are from a multi-gene family encoding cysteine-rich peptides (CRPs), more specifically, a subfamily called defensin-like (DEFL) peptides bearing a cysteine-stabilized alpha-beta motif, as well as a gamma-core (Kanaoka *et al*., 2011; Okuda *et al*., 2009; Takeuchi and Higashiyama, 2012). Further research on these peptides has led to the exciting discovery of two sets of LURE receptors in *Arabidopsis* (Takeuchi and Higashiyama, 2016; Wang *et al*., 2016), as well as the resolution of the structure of one ligand-receptor pair by X-ray crystallography (Zhang *et al*., 2017). Unlike species-specific ZmEA1 and LUREs (Higashiyama *et al*., 2006; Kanaoka *et al*., 2011; Marton *et al*., 2012; Takeuchi and Higashiyama, 2012), which are thought to create a reproductive barrier at the prezygotic level that prioritizes pollen tubes from the same species (Marton *et al*., 2012; Takeuchi and Higashiyama, 2012), XIUQIUs are non-species-specific attractants that increase the overall reproductive success for both conspecific or heterospecific fertilization (Zhong *et al*., 2019). Structurally, except for ZmEA1 in maize (monocot), all known micropylar attractants (LUREs and XIUQIUs) are CRPs.

Our studies use the diploid wild potato species *Solanum chacoense* as a model. Among the 196 wild potato species, *S. chacoense* clearly stands out because of its superior genetic resistance to diseases and pests (Cappadocia, 1990). It has been extensively used in potato breeding through introgression into cultivated tetraploid potato (*Solanum tuberosum*). It is common to have interspecific breeding barriers between wild and cultivated potato, due to a difference in endosperm balance number (EBN, or effective ploidy) of the parents. Such breeding barriers can be overcome by ploidy manipulation and bridge crosses (Jansky, 2006). However, not all crosses can be explained by the EBN theory, and it has been long suggested that other interspecific breeding barriers also exist, acting at the prezygotic and postzygotic levels (Masuelli and Camadro, 1997). It is thus an open question if wild and cultivated potato flowers possess species-specific attractants that form a breeding barrier in addition to the EBN.

In the current study, we first set out to test ovular pollen tube guidance in *S. chacoense* in general by assaying the species-specificity and the impact of glycosylation in pollen tube guidance. We next tried to identify ovule-derived pollen tube attractants in *S. chacoense* and to evaluate their possible contribution to the extensive reproductive isolation in wild and cultivated potato. The identification of candidate attractants was based on transcriptomic comparisons between *S. chacoense* ovules dissected from anthesis flowers (which attract pollen tubes), immature ovules (dissected from flowers at two days before anthesis, 2DBA, which do not attract) and mutant ovules devoid of an embryo sac due to overexpression of *FERTILIZATION-RELATED KINASE 1* (*FRK1*) (which also do not attract) (Lafleur *et al*., 2015; Liu *et al*., 2015). We hypothesized that the 2DBA and mutant ovules do not express pollen tube attractant mRNAs while anthesis ovules do. We performed RNA-seq on these three ovule conditions together with leaf samples. Among the genes identified by differential gene expression (DGE) analyses were 17 ScCRPs which we consider as candidate attractants. These ScCRPs all have a signal peptide, are highly represented in mature ovules, and are under-represented or absent in both immature and *frk1* ovules (2-fold cut-off) and leaf (5-fold cut-off). Based on the patterns of cysteine residues, they can be categorized into 7 CRP subgroups, and the classification, ortholog analyses and alignments of the 17 ScCRPs are described. To evaluate their potential as pollen tube attractants in *S. chacoense*, three DEFL *ScCRPs* were cloned and expressed in a bacterial system. Recombinant peptides showed moderate pollen tube attraction in functional assays (bead assay).

## Material and Methods

### RNA sequencing

*S. chacoense* Bitt. individuals were greenhouse grown at ∼25°C under long-day conditions (16 h light/8 h dark). Flowers were collected from wild type *S. chacoense* at both anthesis (A) and two days before anthesis (2DBA), as well as from a *frk1*mutant line lacking an embryo sac. Flowers at anthesis were also collected from two wild relatives *S. gandarillasii* and *S. tarijense*. Ovaries were hand dissected to remove the pericarp and flash frozen in liquid nitrogen upon harvest. Total RNAs were extracted using TRIzol (Invitrogen).

For transcriptome assembly, two RNA-seq experiments were performed using two different next-generation sequencing platforms as described previously (Liu *et al*., 2015). Briefly, the 454 GS-FLX Titanium platform was first used to sequence the three ovule conditions of *S. chacoense* (A, 2DBA*, frk1*), with each sample occupying one sequencing plate. All cDNA libraries were constructed using a library construction kit (Roche) using an RNA sample enriched for mRNA on oligo(dT)_25_ dynabeads (Invitrogen). Next, an Illumina HiSeq 2000 platform was used to sequence leaf mRNA as well as anthesis stage ovules from *S. chacoense*, *S. gandarillasii* and *S. tarijense,* with each sample occupying one lane. A TruSeq cDNA preparation kit (Illumina) was used to construct the cDNA libraries.

For differential gene expression (DGE) analyses, RNA-seq was performed using Illumina Hiseq on three biological replicates of A, 2DBA, *frk1* ovules and wild type leaves, with each replicate occupying 1/3th of a lane. cDNA libraries were constructed using the TruSeq cDNA preparation kit (Illumina).

### Transcriptome assembly and curation

For *S. chacoense*, 454 reads were assembled into 53,524 contigs using the Newbler package (Margulies *et al*., 2005). Illumina reads were assembled into 47,138 contigs using Trinity (Grabherr *et al*., 2011). A curation protocol was applied to these two primary assemblies. Briefly, contigs from both assemblies were concatenated and first blasted against the NCBI’s refseq_rna database using BLASTn to detect and split chimeric sequences into parts. Resulting sequences were then blasted against the NCBI’s refseq_protein using BLASTp. Frameshifted contigs were resolved by addition of one or two Ns at the appropriate position in the sequence. Due to the large volume of RNA-seq data, contig groups were then generated based on all-versus-all BLASTn comparison. Contigs displaying high level of similarity (≥ 95% sequence identity over ≥ 80% sequence coverage) were grouped and aligned with the MAFFT program (Katoh and Standley, 2013). A consensus sequence was extracted from each alignment to generate a hybrid assembly, on which we performed the above-mentioned chimera and frameshift curation steps again to obtain a final dataset of 45,211 consensus contigs. For *S. gandarillasii* and *S. tarijense*, Illumina reads were assembled using Trinity into 62,171 and 70,188 contigs, respectively.

### Quantification of RNA expression

The expression level of contigs was assessed by mapping reads to the hybrid assembly with Bowtie (Langmead *et al*., 2009). The RSEM program was used to estimate the abundance of contigs and to generate a matrix of raw read counts (Li and Dewey, 2011). The edgeR package was used to compute fold-changes between conditions and the corresponding adjusted *p*-values (Robinson *et al*., 2010).

Semi-quantitative RT-PCR was performed on selected genes using 70 ng of total RNA from anthesis, 2DBA, *frk1* ovules and leaf in biological triplicates using the M-MLV reverse transcriptase kit (Invitrogen). The cDNA templates were amplified for 26 cycles for all genes tested. Ubiquitin was used as the internal control.

### Aniline Blue Staining

*S. chacoense* flowers (G4 genotype, containing S_12_S_14_ self-incompatibility alleles) were pollinated with compatible pollen (V22 genotype, containing S_11_S_13_ self-incompatibility alleles). Forty hours after pollination (by which time fertilization has already occurred), pistils were fixed, squashed and stained as described previously (Matton *et al*., 1997). The pericarps were carefully removed before the pistils were squashed. The pollen tube behavior near the ovule was observed with an Axio Imager M1 microscope equipped with an Axiocam HRc camera (Zeiss).

### Scanning electron microscopy (SEM)

Fertilized pistils were fixed, dehydrated and critical-point-dried as described previously (Gray-Mitsumune *et al*., 2006). Pistils were mounted on double-taped SEM stubs. The pericarps were dissected to reveal pollen tubes prior to coating. Pictures were acquired with a JEOL JSM-35 SEM.

### Protein extraction

*S. chacoense* flowers were collected from the greenhouse. The pericarps were manually removed and the ovules were stored at −80 °C until use. For each extraction, 300 mg of ovules were used. For total protein extraction, samples were ground on ice with a mortar and pestle in liquid nitrogen until a fine powder was obtained. The powder was resuspended in a buffer containing 10 mM Tris·HCl (pH 7.2), 0.5 mM L-cysteine and 0.01% (v/v) Triton X-100. The cold ovule paste was transferred to a 1.5 ml tube and ground again with a plastic pestle on ice until an aqueous extract was obtained. The homogenate was centrifuged for 15 min at 13,000 g at 4 °C. The supernatant was flash-frozen and stored at −80 °C until use. For soluble protein extraction, Triton X-100 was omitted from the buffer. Con A Sepharose 4B beads (GE Healthcare) were used for glycoprotein purification. Here, frozen ovules were ground on ice and total proteins were extracted in ConA binding buffer, containing 10 mM Tris·HCl (pH=7.2), 0.5 M NaCl and 0.5 mM L-cysteine. Five hundred μL bed volume of ConA beads were packed into a column and equilibrated with 10 mM Tris·HCl (pH = 7.2). After loading the total protein extracts, the beads were washed with 10 mM Tris·HCl (pH = 7.2) and eluted with wash buffer containing 1 M methyl-glucopyranoside. Eluents were dialyzed against 10 mM Tris·HCl (pH = 7.2) Before Use in pollen tube attraction assays.

### Expression and Purification of Recombinant Peptides in E. coli

Three *ScCRP* genes belonging to the DEFL family were chosen for *in vitro* pollen tube attraction assays and cloned into the expression vector pET28b. Recombinant peptides were expressed individually in *E. coli* Rosetta-gami 2 and purified under native conditions (without further refolding) using the ÄKTA FPLC system following the instructions for the HisTrap FF 1 mL column. Purified peptides were then concentrated and buffer changed to 50 mM Tris·HCl (pH = 7.8) using an Amicon 3 kDa centrifugal device. Purified peptides were quantified with the Bradford method.

### Pollen tube attraction assays

The single-choice semi-*in vivo* (SIV) assay was performed as described previously (Lafleur *et al*., 2015) using crosses between G4 (♀) x V22 (♂) (Matton *et al*., 1999). In brief, at 24 hours after pollination, full length styles were excised and placed on a BK solid medium (containing 5% sucrose) (Brewbaker and Kwack, 1963). A fresh ovule cluster was dissected from different *Solanum* species and placed at a distance of 700 μm away from the end of the style, at a 45° angle. Pollen tubes exit the style at about 30 hours after pollination. The turning angle of each pollen tube was measured with ImageJ (http://imagej.nih.gov/ij). Angle distribution was compared between conditions using a Kolmogorov – Smirnov (KS) statistical test (*n* > 100).

The two-choice SIV assay was performed as described previously (Lafleur, 2009). Briefly, 24 hours after pollination, full length styles were excised and placed on a BK solid medium (before pollen tubes exit the style) and the protein extracts (1 μL) spotted on the agarose medium using a thin glass capillary inserted into a 0.5-10 μL pipet tip. The protein extraction buffer (1 μL) was used as negative control and placed opposite to the protein extract. Drop applications were performed under a stereomicroscope. In a two-choice SIV assay (involving one style), if pollen tubes are predominately attracted by a protein extract, this assay is marked as “toward extract”. Otherwise, the assay is marked as “toward buffer” or “neutral”. To test for each protein extract, approximately 150 assays were performed. A Student’s *t*-test was used to evaluate the significance of pollen tube attraction between “toward extract” and “toward buffer” populations, based on 4 replicates each. Pollen tube behavior in response to each protein extract was observed in bright field with a Discovery V12 stereomicroscope equipped with an AxioCam HRc camera (Zeiss).

The bead assay was performed following a protocol developed by Okuda, Tsutsui *et al*. (Okuda *et al*., 2009). *S. chacoense* crosses between genotypes V26 (♀) x V22 (♂) were used. To ensure longer pollen tube growth, a full length V26 style was pollinated, then cut at 2/3 its length with surgical scissors, and placed immediately on BK solid medium (containing 12% sucrose and 1.5% low melting agarose). At ∼ 19 hours after pollination, pollen tubes start to exit onto the medium. Gelatin beads supplied with recombinant peptides and 1mM Alexa Fluor dye (Invitrogen) were freshly prepared and assays were begun when pollen tubes were nicely spread out on the medium (∼ 7 hours on the medium). The angle of each pollen tube that appears to be attracted by a gelatin bead was measured by ImageJ.

### Ortholog identification

Orthologs in the genome of *S. tuberosum* and *S. lycopersicum* were retrieved using the NCBI genome data viewer. Orthologs in the transcriptome of *S. gandarillasii* and *S. tarijense* were retrieved using Geneious Prime 2023.2.1 software (https://www.geneious.com). The hits with at least 80% of sequence coverage to the query sequence were manually examined for the presence of a signal peptide and a stop codon. Only the orthologs identified using a reciprocal best BLAST hit method were retained.

### Estimation of dN/dS ratio

Pairwise alignments of the nucleotide sequences encoding the mature peptides were generated with Geneious software using the default parameters. Adjustment was done manually to ensure codon-by-codon alignment. The yn00 program from the PAML package was run to compute pairwise *dN/dS* ratios (Yang and Nielsen, 2000). *Bioinformatic predictions*

Bioinformatic predictions were performed for all transcripts obtained from transcriptome subtraction. Gene Ontology (GO) annotations were obtained using Blast2GO (Conesa and Gotz, 2008). Pfam motif searches were performed using Pfam database (Mistry *et al*., 2021). Protein secretion and glycosylation status was predicted using SignalP 5.0 and NetNGlyc-1.0, respectively.

## Results

### A glimpse at pollen tube guidance in S. chacoense

During a previous study on the collecting method and profiling of ovule secretome in *S. chacoense*, we observed pollen tubes are attracted to ovule clusters from the same species (Liu *et al*., 2015). In this work, we then went further to ask (1) whether this attraction remains when ovules from the same species are replaced by ovules from closely related or distant *Solanum* species; (2) whether post translational modification (glycosylation) has an impact on the attraction. These questions thus address two different characteristics of pollen tube guidance in this species.

### S. chacoense pollen tubes show species-specificity toward ovules of their own species

In *Arabidopsis thaliana*, two sets of micropylar attractants have been discovered: LURE peptides that are species-specific and direct pollen tubes to fertilize ovules of its own species (Higashiyama *et al*., 2006; Kanaoka *et al*., 2011; Takeuchi and Higashiyama, 2012) ; and XIUQIU peptides that are non-species-specific (Brassicaceae-conserved) and increase the overall reproductive success for both conspecific or heterospecific fertilization (Zhong *et al*., 2019) .

Compared to *Arabidopsis*, does pollen tube attraction show species-specificity in wild potato? Pollen tube attraction to several *Solanum* species was previously examined using a single-choice SIV assay (Lafleur, 2009). In this study, pollen tubes emerging from the style could go toward the ovule cluster side (attraction), toward the opposite side (no attraction), or grow into a dome-like shape without showing a preference (neutral). The percentage of assays (styles) showing attraction were recorded for a total of 8 *Solanum* species crosses. The results showed that, in 47 out of 80 pollinated *S. chacoense* styles, pollen tubes were predominately attracted by ovules of the same species (reported as 59% (*n* = 80)) (Lafleur, 2009). In contrast, *S. chacoense* pollen tubes were attracted to ovules from distant species at a much lower rate. Specifically, *S. chacoense* pollen tubes are attracted to *S. bulbocastanum* ovules at 22% (*n* = 55); to *S. tarijense* ovules at 21% (*n* = 48); to *S. pinnatisectum* ovules at 14% (*n* = 43); to *S. microdontum* ovules at 11% (*n* = 71), to *S. commersonii* ovules at 10% (*n* = 52); to *S. tuberosum* ovules at 9% (*n* = 45), and to *S. lycopersicum* ovules at 0% (*n* = 55). *S. chacoense* pollen tube attraction toward ovules from all other *Solanum* species was significantly different from that of its own species according to a Chi-squared test (*p* < 0.0001), indicating *S. chacoense* pollen tubes were preferentially attracted to ovules from their own species.

Here, we have repeated these SIV assays with *S. tarijense* in order to compare them to S. *gandarillasii*. While both S. *gandarillasii* and *S. tarijense* are closely related to *S. chacoense*, *S. gandarillasii* is phylogenetically closer to *S. chacoense* than *S. tarijense* (Rodríguez and Spooner, 2009). In these assays, the cumulative distribution of each pollen tube curvature growing towards ovules from two species was compared using Kolmogorov–Smirnov (KS) test. Fig 1A – C shows typical pollen tube behavior towards ovules from *S. chacoense*, *S. gandarillasii* and *S. tarijense*. When *S. chacoense* pollen tubes were given a choice of *S. gandarillasii* ovules and *S. chacoense* ovules, pollen tube behavior near *S. gandarillasii* ovules was not significantly different from that of *S. chacoense* ovules (both attracting pollen tubes equally, *n* > 100). In contrast, *S. tarijense* ovules induced significantly different pollen tube behavior than that of *S. chacoense* (*n* > 100, *p* < 0.001), rejecting the null hypothesis that the two conditions are drawn from the same distribution.

**Fig 1.**
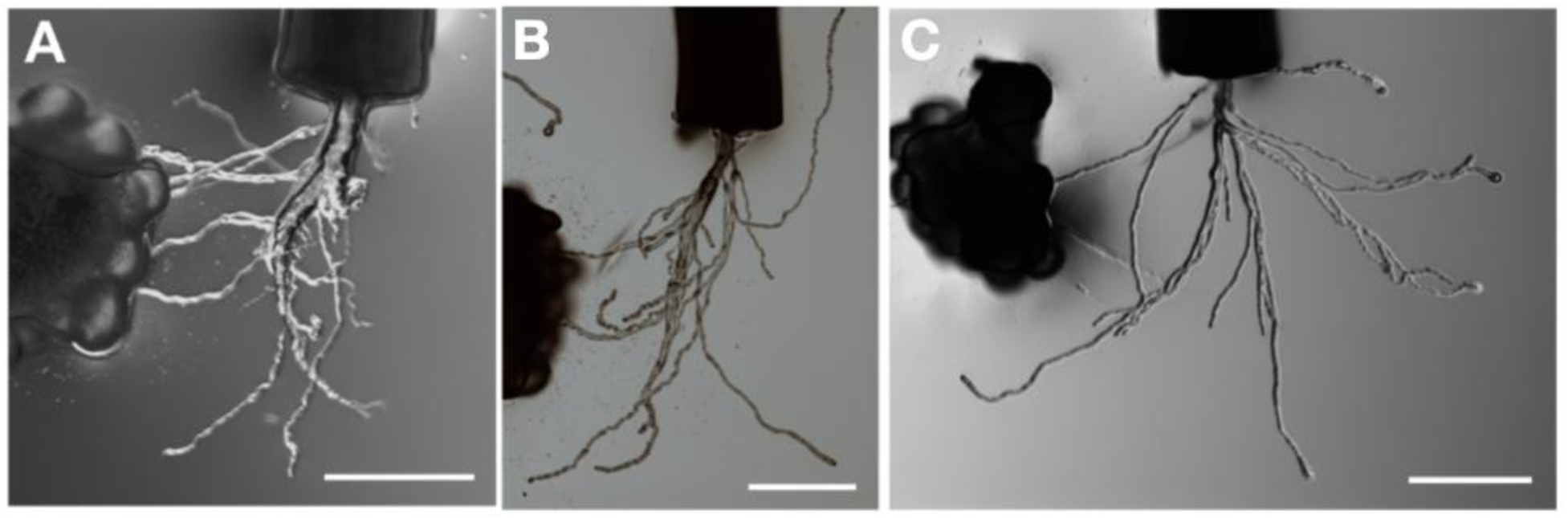
Chemotropism of *S. chacoense* pollen tubes toward different ovule sources in the single-choice SIV assay. (A) toward *S. chacoense* ovule clusters, (B) toward *S. gandarillasii* ovule clusters and (C) toward *S. tarijense* ovule clusters. Scale bar = 400 μm.

Thus, based on our SIV assays of close relatives, combined with previous SIV assays using more distantly related *Solanum* species (Lafleur, 2009), we conclude that pollen tube attraction functions in a species-specific manner in wild potato species, as has been reported in *Torenia*, *Arabidopsis* and maize (Kanaoka *et al*., 2011; Marton *et al*., 2012; Takeuchi and Higashiyama, 2012). Furthermore, our results also suggest a positive correlation between species-specificity of pollen tube guidance and their taxonomic distance, as *S. chacoense* pollen tubes are attracted to both *S. chacoense* and S. *gandarillasii* ovules more than to *S. tarijense* ovules.

### The glycoprotein fraction of total ovule extracts enhances pollen tube attraction

Glycosylation of plant proteins is known to affect quality control in the endoplasmic reticulum, protein stability, and ligand receptor interactions among others (Strasser, 2016). N-glycosylation in particular is a common post translational modification for proteins entering the secretory pathway. To assess if glycosylation impacted pollen tube guidance in *S. chacoense*, the two-choice assay system described previously (Lafleur, 2009) was used to evaluate three protein extracts from ovules at anthesis on pollen tube attraction. These include a total protein extract (extraction buffer contains detergent), a soluble protein fraction (extraction buffer lacks detergent) and a glycoprotein fraction (prepared from the soluble protein fraction).

**Fig 2.**
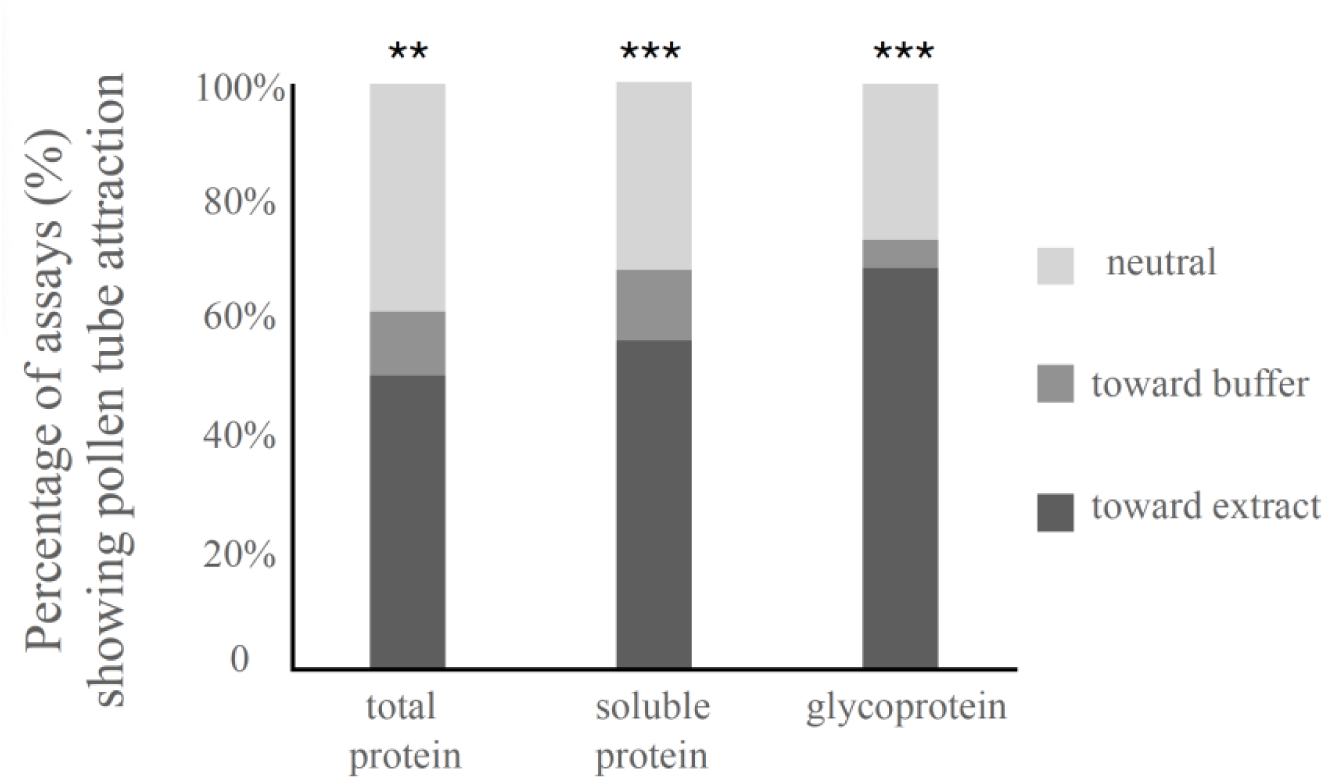
Summary of the percentage of assays showing pollen tube attraction in the two-choice assay system. Toward extract category sums the total number of styles that have the majority of pollen tubes grow toward protein extracts. Neutral represents styles where the majority of pollen tubes grow symmetrically toward both sides. The *p*-values, calculated by Student’s *t*-test between “toward extract” and “toward buffer” populations, were based on four replicates for each protein extract (\**р* < 0.05; \*\**p* < 0.01; \*\*\**р* < 0.001).

As shown in Fig 2, when total protein extracts are placed on one side of pollen tubes exiting from the style, in 76 out of 151 assays (involving 151 styles) pollen tubes grow towards total protein extracts (*n* = 151, *p* < 0.05). When total protein extract was replaced by soluble and glycoprotein extracts, this percentage increased to 56% (*n* = 149, *p* < 0.001) and 68% (*n* = 148, *p* < 0.001), respectively. The glycoprotein fraction of total ovule extracts thus increased the number of experiments where pollen tubes were attracted from half to two-thirds, suggesting this post-translational modification may be important to the process of pollen tube guidance in *S. chacoense*. Typical images of pollen tube attraction by glycoprotein extracts are shown in Fig 3.

**Fig 3.**
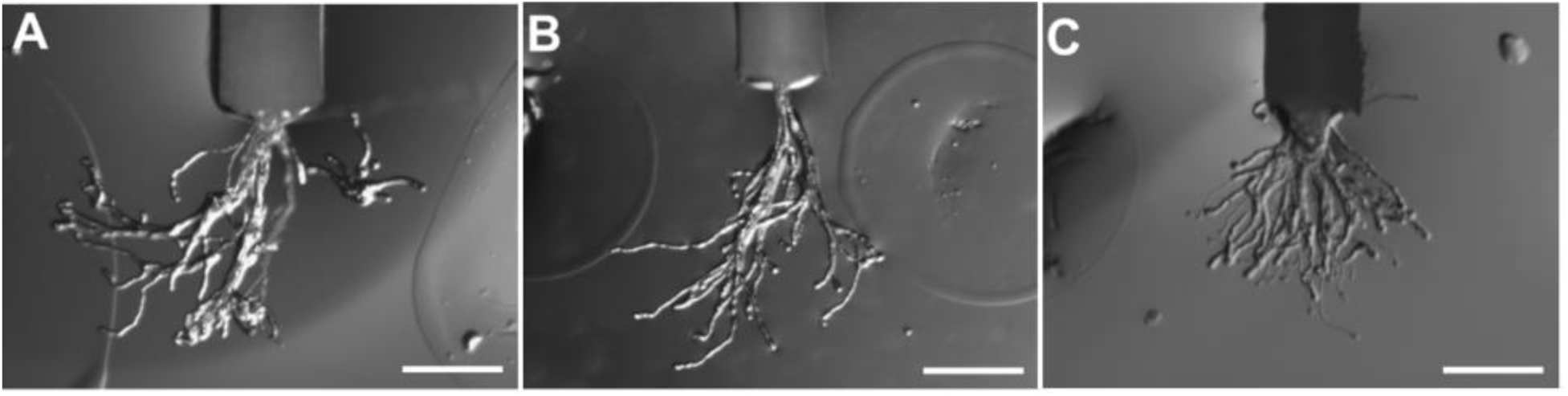
A glycoprotein fraction from total ovule extracts attracts pollen tubes in the two-choice assay system. (A, B) A glycoprotein fraction was deposited on the left side of the style while the extraction buffer used during glycoprotein purification was deposited on the right side, roughly equidistant from the pollen tubes exiting the style. The drops left a circular print on each side of the style. (C) Extraction buffer was deposited on both sides of the style. Scale bar = 400 μm.

### frk1 mutant ovules show defects in pollen tube guidance in vivo

We have previously shown that neither immature ovules dissected at two days before anthesis flowers (2DBA) or *frk1* mutant ovules (devoid of an embryo sac) attract growing pollen tubes in a SIV assay (Lafleur *et al*., 2015; Liu *et al*., 2015). To further examine the phenotype of *frk1* mutants regarding pollen tube guidance in *vivo*, we hand-pollinated two self-incompatible but cross compatible genotypes of *S. chacoense* and visualized pollen tube behavior inside the ovary by aniline blue staining and SEM imaging. Results show contrasting pollen tube behavior near wild-type versus *frk1* ovules (Fig 4A and 4B). In Fig 4A, as multiple pollen tubes approached wild-type mature ovules, one or several pollen tubes turned sharply toward each ovule. This pattern was not seen using *frk1* plants, where pollen tubes grew randomly around the ovules (Fig 4B). SEM images provide better visualization of individual pollen tubes and ovules (Fig 4C and 4D). As shown in Fig 4C, all pollen tubes migrated consistently on the surface of the placental tissue of wild-type ovules and no pollen tubes were seen on the surface of the ovules themselves. The micropyle of *S. chacoense* ovules faces towards the placenta and is deeply hidden under nucellus tissues in these images so visualization of pollen tubes entering the micropyle is not possible. In contrast to the wild type, multiple pollen tubes migrated randomly on the surface of *frk1* ovules, demonstrating a defect in pollen tube guidance *in vivo*.

**Fig 4.**
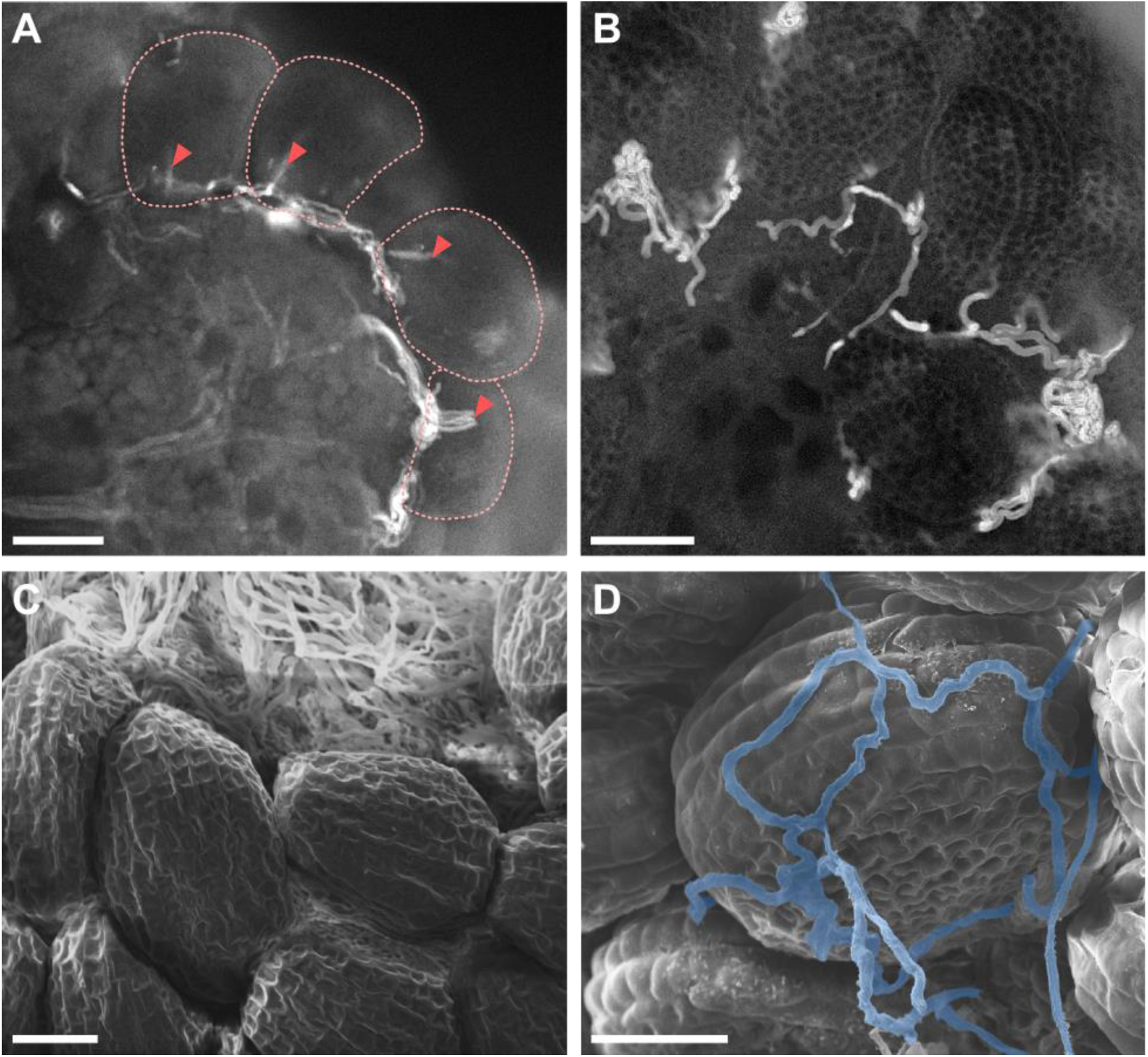
*in vivo* pollen tube guidance by wild-type and *frk1* ovaries visualized by aniline blue staining (A, B) and SEM (C, D). (A) Pollen tubes targeting the micropyle of wild-type ovules. Dotted red lines indicate the contour of each single ovule. Red arrowheads point at the pollen tubes that turned sharply to target the micropyle for double fertilization. (B) Pollen tubes migrating randomly on the surface of *frk1* ovules. (C) Pollen tube behavior near wild-type ovules showing pollen tubes elongated along the placental epithelium. (D) Pollen tube behavior near *frk1* ovules. Pollen tubes that elongated randomly on the surface of the ovule were colored in blue. Scale bar = 100 μm.

### DGE identifies 17 CRP candidate attractants for ovular pollen tube guidance in S. chacoense

#### Selection criteria

DGE analyses were performed using the *S. chacoense* transcriptome assembly. The goal was to identify genes that are highly expressed in ovules able to attract pollen tubes while having low/zero expression in ovules that are unable to attract pollen tubes. In all, three different ovule samples and one leaf sample were compared. The ovule samples were 1) mature ovules from anthesis flowers that attract pollen tubes (called **A**); 2) immature ovules from flowers at two days before anthesis (**2DBA**) that do not attract pollen tubes; 3) ovules from a *frk1* mutant that lack an embryo sac and also do not attract pollen tubes. The leaf tissue was used as sporophytic control that never produces ovular attractants.

The following selection criteria used to select transcripts encoding potential attractants were, 1) presence of a signal peptide to allow secretion; 2) abundant expression in mature ovules that was two-fold greater than in both 2DBA and *frk1* ovules (*p* < 0.01) and five-fold greater than in leaf samples (*p* < 0.01). A total of 284 transcripts fulfilled these criteria, which are thus considered as candidates for ovular pollen tube attractants in *S. chacoense*. These transcripts are provided in Supplementary Table S1, along with their annotations, expression and fold change information.

To date, all known micropylar attractants (such as LUREs and XIUQIUs in *Torenia* and *Arabidopsis*) are CRPs. We thus decided to focus our attention on CRPs in the gene list of potential attractants. The search for CRPs used KAPPA, an algorithm designed for discovery and clustering of proteins with a set pattern of cysteine residues (Joly and Matton, 2015). A subset of 17 CRPs bearing at least 4 cysteines in the protein backbone was retained.

Before examining the 17 CRPs more closely, RT-PCR was performed as a means to validate DGE prediction. As shown in Supplementary Fig. S1, RT-PCR showed a unique expression in anthesis ovules for the majority of the *ScCRP* genes, as opposed to low or zero expression in 2DBA, *frk1* ovules and leaf. RT-PCR results are consistent with DGE data and thus provide confidence in our candidate selection strategy. These genes are now termed *ScCRP*s. The primers used for RT-PCRs are listed in Supplementary Table S2.

The length of the mature CRP sequences vary between 54 to 192 amino acids (aa). Of these, 10 genes encode mature peptides under 100 aa, 6 genes encode mature proteins between 100 – 150 aa, while 1 gene encodes a mature protein of 192 aa. Typically, mature CRPs are up to 100 aa in length. All 17 sequences were included in the following analyses, despite some being longer than previously identified CRPs. Four of the ScCRPs had only a partial sequence in the transcriptome, and their sequences were completed using a previously published *S. chacoense* genome (Leisner *et al*., 2018). The coding sequence and protein sequences, as well as DGE data and fold changes between different ovule conditions, are provided in Supplementary Table S3.

#### CRP Classification

To determine if any known function might be associated with our 17 ScCRPs, an initial protein BLAST was done against the non-redundant database in NCBI using default parameters. A previous study (Silverstein *et al*., 2007) examined the cysteine arrangement of expressed sequences across 33 plant species and categorized all plant CRPs into 516 subgroups. A second protein BLAST was thus done against this plant CRP list in NCBI to place our CRPs into the known subgroups. The classification was validated using sequence alignments between each CRP and selected example sequences in the corresponding CRP subgroup using Geneious software, with manual correction on certain regions. The alignments are shown in Supplementary Fig. S2, showing the location of signal peptide as well as the cysteine arrangement in each CRP subgroup.

This analysis indicated that 7 candidate CRPs (mature length between 103 – 192 aa) belong to the early culture abundant 1 (ECA1) gametogenesis-related protein family, 4 belong to the defensin-like (DEFLs) protein family (mature size between 54 – 60 aa), and 6 (mature length between 49 – 81 aa) do not resemble any previously described CRP subgroups and are thus classified as novel CRP families. Candidate CRP classification is summarized in Table 1 and Supplementary Table S3.

**Table 1.**
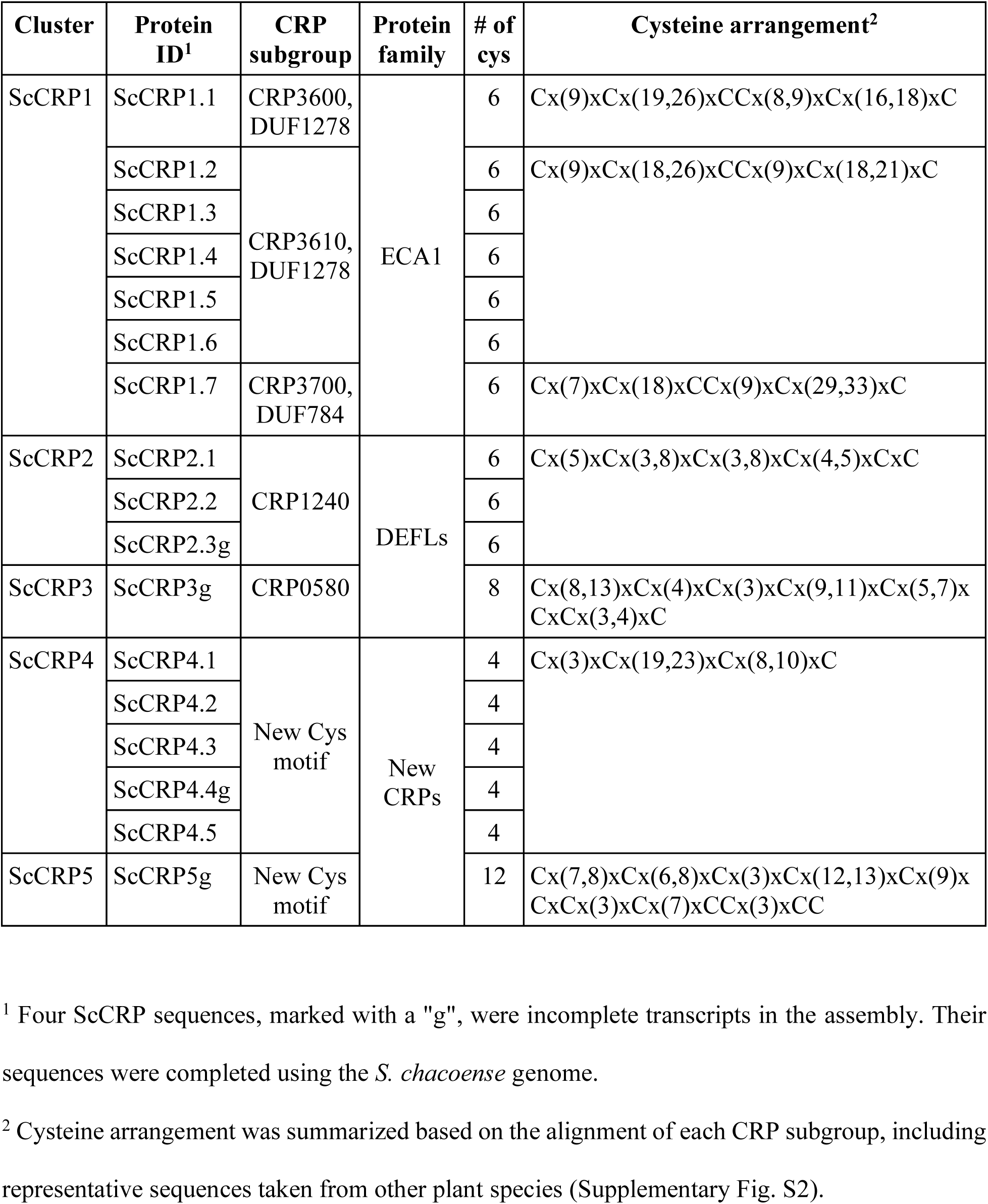
Candidate ScCRP classification.

### ScCRP1 members belong to the ECA1 gametogenesis-related protein family

The ECA1 gametogenesis-related protein family is largely unexplored. Its members are characterized by the prolamin-like domain containing six cysteines spaced as C1…C2…C3C4…C5…C6. Three disulfide bonds (C1-C5, C2-C3, and C4-C6) have been suggested (Sprunck *et al*., 2014), and the disulfide bond formation of ScCRP1.1-1.6 predicted by AlphaFold 3 is in agreement with this arrangement. Based on different spacing between cysteine residues, several CRP subgroups have been identified and listed between CRP 3600 and CRP 3740 (Silverstein *et al*., 2007). Our sequence alignment indicates that ScCRP1.1 belongs to the CRP3600 subgroup, ScCRP1.2–1.6 belong to the CRP3610 subgroup, and ScCRP1.7 belongs to the CRP3700 subgroup.

Interestingly, a known function has been associated with the CRP3610 subgroup (containing ScCRP1.2–1.6). This subgroup contains five orthologous *EC1* (egg cell 1) genes that have been well studied in *Arabidopsis thaliana* (Sprunck *et al*., 2012). EC1 interacts with and activates sperm cells for gamete fusion right before double fertilization. When aligned with EC1 peptides, ScCRP1.2–1.6 share the two common EC1 signature motifs S1 and S2, as well as a conserved tryptophan residue following the fifth cysteine residue (Supplementary Fig. S2).

### ScCRP2 and ScCRP3 belong to the DEFL protein family

ScCRP2.1–2.3 all belong to a multi-gene family in the subgroup CRP1240, all sharing a common 6 cysteine motif. The mature peptide length in this subgroup ranges between 54 and 60 amino acids, with percentage of identity in their alignment ranging between 39.5% and 44.7%. When aligned with example sequences from this subgroup (Supplementary Fig. S2), a common cysteine arrangement can be summarized as: Cx(5)xCx(3,8)xCx(3,8)xCx(4,5)xCxC.

ScCRP3 also belongs to the DEFL family but is in the CRP0580 subgroup. It has an 8-cysteine motif in its mature peptide of 58 amino acids with a cysteine arrangement of Cx(8,13)xCx(4)xCx(3)xCx(9,11)xCx(5,7)xCxCx(3,4)xC.

### ScCRP4 and ScCRP5 are novel CRPs discovered in S. chacoense

ScCRPs4.1–4.5 did not fit into any of the CRP subgroups reported by Silverstein’s group and did not show similarity to other known proteins in the nr database. We report them here as novel CRPs in *S. chacoense* ovules. A common 4 cysteine motif was identified with cysteine arrangement summarized as: Cx(3)xCx(19,23)xCx(8,10)xC. The mature peptide length in this multi-gene family ranges between 49 and 62 amino acids. When aligned, the percentage of identity within its members range between 33.3% and 71.4%.

ScCRP5, bearing a larger number of cysteines (12-cysteine motif), is also classified as a novel CRP in *S. chacoense* ovules. The ScCRP5 transcript was incomplete from our transcriptome assembly and was completed using the *S. chacoense* genome. The cysteine arrangement is: Cx(7,8)xCx(6,8)xCx(3)xCx(12,13)xCx(9)xCxCx(3)xCx(7)xCCx(3)xCC. Although ScCRP5 did not fit into any known CRP subgroups, BLAST hits from *Nicotiana* species were found in the nr database. The hits show sequence identity to ScCRP5 between 21.4% and 33.3%. However, they only align with the first part of ScCRP5.

### dN/dS ratios indicate 4 ScCRPs belonging to the DEFL and novel CRP families may undergo positive selection

The species-specific pollen tube attraction in *Solanum* noted above suggests that attractants secreted from the ovules of one species may undergo positive selection (changing the amino acid sequence of its ortholog) in different species in order to promote conspecific pollen tube attraction. The *dN/dS* ratio of the orthologs of our 17 ScCRPs were thus investigated. The genome sequences of *S. tuberosum* (The Potato Genome Sequencing Consortium, 2011) and *S. lycopersicum* (The Tomato Genome Consortium, 2012) were both searched for orthologs. In addition, orthologs were also sought in our *S. tarijense* and *S. gandarillassi* ovule transcriptomes. Orthologs were found in at least two relatives of *S. chacoense* using the reciprocal best BLAST hit method (Supplementary Table S4).

To evaluate the selection pressures acting on CRP protein coding regions, the rate of substitutions at non-silent sites (*dN*; non-synonymous substitution) versus substitutions at silent sites (*dS*; synonymous substitutions) were calculated for each ScCRP compared to its orthologs. A *dN/dS* ratio larger than 1 suggests natural selection promotes changes in the protein sequence (positive selection), while a *dN/dS* ratio less than 1 suggests natural selection suppresses protein changes (negative selection) (Kryazhimskiy and Plotkin, 2008). As shown (Supplementary Table S5), ScCRP1 cluster proteins (ScCRP1.1–1.7), which belong to the ECA1 family, all have *dN/dS* ratio less than 1 in pairwise comparison to other *Solanum* species indicating that their protein sequences are conserved among different *Solanum* species. However, for ScCRPs belonging to DEFL and novel CRP families, out of the 23 orthologs found in other *Solanum* species, 8 pairs have a *dN/dS* ratio > 1. These pairs (ScCRP2.1/SgCRP2.1, ScCRP2.1/StCRP2.1, ScCRP2.1/StubCRP2.1, ScCRP3/StubCRP3, ScCRP3/SlCRP3, ScCRP4.2/SlCRP4.2, ScCRP4.4/SgCRP4.4 and ScCRP4.4/StCRP4.4) suggest that natural selection promotes more protein sequence changes in ScCRP2.1 (DEFL), ScCRP3 (DEFL), ScCRP4.2 and ScCRP4.4 (novel CRPs).

### Recombinant ScCRP2.1–2.3 peptides (DEFLs) attract growing pollen tubes

The known attractants, *TfLUREs* and *AtLUREs*, both belong to a multigene family encoding DEFL peptides. Out of the 17 *ScCRPs* identified by our transcriptomic approach, *ScCRP2.1; 2.2* and *2.3* seem to fit in this category. We thus tested them for *in vitro* pollen tube attraction activity using a bead assay, a design that allows individual pollen tube to be tested for attraction to proteins embedded in gelatin beads (Okuda *et al*., 2009). To do so, the three genes were cloned, expressed, and purified from soluble fractions under native conditions without further refolding.

The identification of purified ScCRP2.1 was confirmed by mass spectrometry sequencing before the functional assays.

We then optimized the bead assay for *S. chacoense* by cutting the style at 2/3 length (with surgical scissors) and increasing sucrose concentration to 12% in the pollen tube growth medium. These changes allowed longer elongation times of pollen tubes on the medium. Recombinant peptides were tested in the bead assay after pollen tubes had grown for about 7 hours on medium, a time frame when *in vitro* pollen tube attraction in *Torenia* was observed (Okuda *et al*., 2013). We observed moderate level of pollen tube attraction activity in all three peptides. Representative images of pollen tube attraction activity are shown in Fig 5A–D.

For ScCRP2.2, 56% of pollen tubes were attracted by 230 μM (*n* = 27), while 23 μM ScCRP2.2 attracted 21% of pollen tubes (*n* = 19). As shown in Fig 5B, we observed multiple pollen tubes approaching and being attracted by ScCRP2.2 simultaneously. The turning angle was recorded for each pollen tube being attracted, and the average angle of attracted pollen tubes for ScCRP2.2 was 50°. In parallel, we found 28% of pollen tubes were attracted by 35 μM ScCRP2.1 (*n* = 29), with an average of turning angle of 55° and 41% of pollen tubes were attracted by 2 μM ScCRP2.3 (*n* = 22), with an average of turning angle of 35°. Only two out of 29 pollen tubes (7%) seemed attracted by buffer alone.

**Fig 5.**
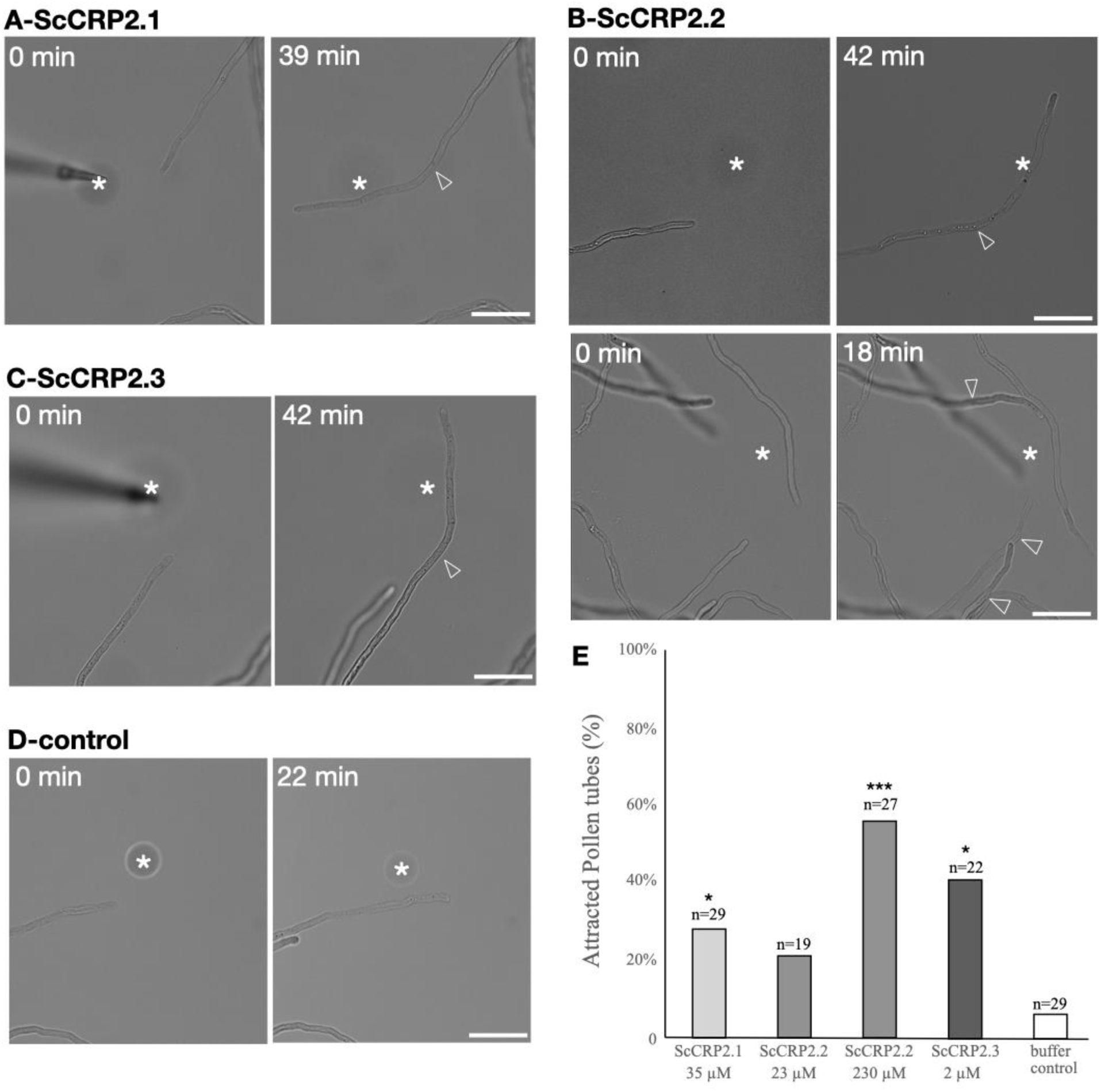
*in vitro* pollen tube attraction assays using recombinant CRPs. (A) A gelatin bead containing ScCRP2.1 recombinant peptides and 10 kD Alexa Fluor 488 was placed on the path of a growing pollen tube through a micropipette. Images were taken at time zero as well as at 39 min showing ScCRP2.1 peptides attracted the pollen tube. Arrowheads mark the position of the tip of the pollen tubes when gelatin beads (asterisks) were placed. Scale bar = 50 μm. (B, C) Representative images show ScCRP2.2 and ScCRP2.3 recombinant peptides attracting pollen tubes. Scale bar = 50 μm. (D) A gelatin bead containing Tris HCl buffer (pH = 7.8) did not attract the pollen tube. Scale bar = 50 μm. (E) Percentage of attracted pollen tubes by recombinant peptides of ScCRP2.1–2.3 at tested concentrations. The data are the percentage for the total number of pollen tubes (*n*) in three or four experiments per peptide using from 4 –12 pollen tubes (except for ScCRP2.2 at 23 μM which has two experiments). Asterisk(s) indicate the significant differences compared with buffer alone using Student’s *t*-test (\**p* < 0.05; \*\**p* <0.01; \*\*\**p* <0.001)

## Discussion

### Transcriptome subtraction of three ovule conditions defines genes involved in late stage of embryo sac maturation

In this study, we used transcriptome subtraction to identify transcripts that were highly expressed in ovules that attract pollen tubes (anthesis, A) while being poorly expressed in ovules that do not (2DBA and in the embryo sac-less mutant *frk1* (Lafleur *et al*., 2015)). A list of 284 genes was obtained as a candidate gene pool for ovular pollen tube guidance attractants in *S. chacoense* (Supplementary Table S1). Among these were found 17 ScCRPs, considered to be prime pollen tube attractant candidates since previously identified micropylar attractants in *Torenia* and *Arabidopsis* are all CRPs (Kanaoka *et al*., 2011; Okuda *et al*., 2009; Takeuchi and Higashiyama, 2012; Zhong *et al*., 2019) (Table 1, Supplementary Table S3).

A more precise description of the developmental stage for each type of ovule used would include that (1) sample A represents wild-type ovules that contain a mature embryo sac at the FG7 stage; (2) sample 2DBA (two days before anthesis) represents slightly immature wild-type ovules (bud size 6–7 mm), which contain a developing embryo sac at the FG6 stage (unfused polar nuclei) while the overall morphology resembles the mature ones (Chevalier *et al*., 2013); (3) sample *frk1* represents embryo sac-devoid mutant ovules either where embryo sac development is stopped at the functional megaspore stage (FG1) or where the embryo sac is entirely lacking. Thus, compared to wild-type mature ovules, the use of *frk1* ovules defines embryo sac-dependent genes while the use of 2DBA ovules further defines a subset of embryo sac-dependent genes that are specifically involved in late stages of embryo sac maturation (the FG6 to FG7 transition). It should be noted that, since the samples sequenced were derived from whole ovules (lacking the ovary wall), genes expressed from the sporophytic tissues of the ovule (nucellus and integuments) should also be expected in the final 284 gene list. However, the expression of all genes in the list depends on the presence of an embryo sac. Detailed information for the 284 gene list, including the basic information of each gene, best informative BLASTp hit from nr database, GO annotations, FPAM annotations, as well as DGE analyses between ovule conditions is provided (Supplementary Table S1).

The gene list contains several genes other than CRPs that are of interest. For example, ScOT_14006 shares 51.4% protein identity to AtMYB108 and 51.7% to SlMYB21, transcription factors known to control stamen development in *Arabidopsis* (Mandaokar and Browse, 2009) and ovule development in tomato (Schubert *et al*., 2019). As another example, ScOT_21090 shares 35.9% protein identity to GPI-anchored LORELEI in *Arabidopsis*, which is localized to the filiform apparatus between two synergid cells (Liu *et al*., 2016) and is known to play a role in pollen tube reception by the female gametophyte (Tsukamoto *et al*., 2010). The list also contains proteins of unknown function ScOT_15973 (DUF639, PF04842) and ScOT_17645 (DUF1442, PF07279), as well as several proteins that have no hits in nr databases. Preliminary analyses for genes potentially involved in embryo sac development, late stage of embryo sac maturation (FG6-FG7), as well as in pollen tube guidance in *S. chacoense* are provided in Supplementary Table S1.

### Are ScCRP candidates associated with long-range guidance, funicular guidance or micropylar guidance?

So far, three types of ovular pollen tube guidance have been distinguished, long-range guidance, funicular guidance and micropylar guidance. Only the attractants in micropylar guidance step have been identified as species-specific ZmEA1 in maize, LUREs in *Torenia* and *Arabidopsis* (Kanaoka *et al*., 2011; Okuda *et al*., 2009; Takeuchi and Higashiyama, 2012) and XIUQIU peptides (Zhong *et al*., 2019) in *Arabidopsis* and Brassicaceae.

Long-range guidance features a transition point when pollen tubes switch from an intercellular growth mode (in the transmitting tract) to surface growth mode (on the septum). The embryo sac appears to direct this long-range signaling process (Hulskamp *et al*., 1995). Funicular guidance, however, has proved difficult to demonstrate. In particular for *S. chacoense*, the morphology of the ovule determined using SEM shows the funiculus to be extremely short, meaning that the micropylar opening of the ovule is in the immediate vicinity of the placental surface, giving direct access to approaching pollen tubes (Liu *et al*., 2015). This type of signaling may be of less importance in this species.

Finally, micropylar attractants (LUREs and XIUQIUs) are produced by the two synergid cells within the embryo sac (Higashiyama *et al*., 2001; Zhong *et al*., 2019). Since the transcriptome subtraction method used here identifies genes whose expression depends on the presence of an embryo sac (Supplementary Table S1), the 17 ScCRPs reported here could thus be associated with either long-range guidance or micropylar guidance.

### Investigation of the 17 ScCRPs provides insights into embryo sac-dependent, small secreted CRPs by S. chacoense ovules

In this research, we identified 17 ScCRPs as prime candidates for ovular pollen tube guidance attractants in the wild potato species *S. chacoense* (Supplementary Table S3). Analysis of their cysteine patterns, classification, alignment, orthologs in other *Solanum* species and the strength of natural selection (measured by *dN/dS* ratio) were all examined to characterize ScCRPs differentially regulated during ovule development and their potential as ovular pollen tube attractants in *S. chacoense*. The selection of the 17 ScCRPs was made using DGE analyses (showing fold changes between three ovule conditions; Supplementary Table S1). Details regarding the examination of their cysteine arrangement accompanied by CRP classification (Table 1), alignments of each CRP subgroup (Supplementary Fig. S2), ortholog analyses in closely related as well as in distant *Solanum* species (Supplementary Table S4), and pairwise *dN/dS* ratio estimations between orthologs (Supplementary Table S5) are provided. Among the 17 ScCRPs are found 7 ScCRPs belonging to ECA1 gametogenesis-related protein family, 4 ScCRPs identified as DEFLs and 6 novel ScCRPs based on the appearance of new cysteine motifs.

*Arabidopsis thaliana* has 745 CRPs, and of these, 124 encode ECA1 family members as determined by SPADA (Zhou *et al*., 2013). In two separate reports (Jones-Rhoades *et al*., 2007; Punwani *et al*., 2007) a combined total of 43 ECA1 family members whose expression depends on the synergic cell-expressed transcription factor *MYB98* (a key regulator in micropylar pollen tube guidance) were discovered in *Arabidopsis thaliana* (Kasahara *et al*., 2005). Using promoter GUS constructs, the subcellular localization of 20 of these 43 ECA1 was investigated, and all show solely or primary expression in the synergic cell of the female gametophyte (Jones-Rhoades *et al*., 2007; Punwani *et al*., 2008). Considering the fact that synergid cells are the main source of pollen tube attractants in *Torenia* and *Arabidopsis*, the ECA1 family candidates ScCRP1.1–1.7 are of particular interest. However, because the function of ECA1 gametogenesis-related protein family members in general is largely unknown, it will be important to examine their involvement in pollen tube guidance. The EC1 peptides whose functions are known in *Arabidopsi*s are orthologous to ScCRP1.2–1.6, and their function is to trigger sperm activation and gamete interaction. EC1 is thus the only member in the ECA1 family that has a defined function. It would be of interest to examine whether ScCRP1.2–1.6 might have a similar function in *Solanum*, or if these proteins take on new roles as attractants in *S. chacoense*.

For the DEFLs, *Arabidopsis* has 317 genes encoding DEFL peptides (Silverstein *et al*., 2005). In addition to the well-known role of LUREs as micropylar attractants in pollen tube guidance, DEFLs have been reported to play other roles, such as in plant defense (Hawamda *et al*., 2022), self-incompatibility in *Brassica* species (Takayama *et al*., 2000), and pollen tube tip bursting in maize (Amien *et al*., 2010). Among our DEFLs candidates, ScCRP2.1–2.3 belong to CRP1240 while ScCRP3 is a singleton belonging to CRP0580. None have any previously reported function.

The novel ScCRPs 4.1–4.5 and ScCRP5, that has cysteine motifs unique to *Solanum* species, will also be of interest to further characterize. It is possible new roles could be assigned to these small peptides, possibly from a wider perspective as in embryo sac development, pollen-pistil interaction and in flowering plant reproduction.

### Functional assays confirm ScCRP2.1–2.3 (DEFLs) are good candidates as ovular attractants in S. chacoense

When ScCRP2.1–2.3 recombinant peptides were tested in functional assays, moderate levels of pollen tube attraction activity were observed for all three. Here we exploited the same bead assay developed and used for the visualization of *in vitro* pollen tube attraction activity in *Torenia* and, later on, in *Arabidopsis*. It was reported that in *Torenia fournieri*, LURE1 and LURE2 show significant pollen tube attraction in a concentration-dependent manner. At the optimal concentration, 52.5 ± 3.4% (*n* = 34) of pollen tubes were attracted by 40 nM LURE1 while 65.7 ± 7.4% (*n* = 44) of pollen tubes were attracted by 4 nM LURE2 (Okuda *et al*., 2009). In *Arabidopsis*, 100% of pollen tubes were attracted by AtLURE1.2 at the optimal concentration of 50 μM (Takeuchi and Higashiyama, 2012). A group of XIUQIU peptides attracted *Arabidopsis thaliana* pollen tubes at 62.2 ± 9.1%, 21.0 ± 7.2%, 3.0 ± 5.2% and 28.3 ± 10.4%, depending on the peptide (Zhong *et al*., 2019).

For *S. chacoense*, ScCRP2.2 was tested at two concentrations (23 μM and 230 μM) similar to those mentioned above. We observed 56% of pollen tubes were attracted by 230 μM ScCRP2.2 (*n* = 27). ScCRP2.1 and ScCRP2.3 were tested at only one concentration. Specifically, 28% pollen tubes were attracted by 35 μM ScCRP2.1 and 41% by 2 μM ScCRP2.3. These attraction activities align well with the LUREs in *Torenia* and the XIUQIU peptides in *Arabidopsis*.

In the bead assay, regular pollen tube growth medium was used instead of special medium pre-conditioned by ovules. In the study of *Torenia* pollen tube guidance, an arabinogalactan polysaccharide termed AMOR was identified that makes pollen tubes competent to respond to LUREs (Mizukami *et al*., 2016). The impact of conditioned medium in pollen tube attraction in *S. chacoense* has not been investigated.

Taken together, the data shown here suggests the multigene family *ScCRP2.1*–*2.3* members are prime candidates that direct pollen tube guidance in *S. chacoense*. Furthermore, pairwise *dN/dS* comparison between *S. chacoense* and its homologs suggest natural selection promotes more protein sequence changes in *ScCRP2.1*, suggesting ScCRP2.1 may be involved in species-specific pollen tube attraction (Supplementary Table S5). Regarding the aspect of glycosylation, six of the 17 ScCRPs (35%) were predicted to contain at least one potential N-glycosylation site. No potential glycosylation site is predicted for ScCRP2.1–2.3 (Supplementary Table S6).

Following *in vitro* assays, it will be important to look into the subcellular localization of these genes, to evaluate their impact on fertilization and seed set through knockout plants, and thus to examine if they are true attractants that direct pollen tube guidance in wild potato species.

## Conclusion

In this report, we examine the pollen tube guidance mechanism in the wild potato *Solanum chacoense*. Our particular interest lies in ovular attractants which act at a stage when pollen tubes are guided to approach and enter the micropylar opening of the ovule before double fertilization. In an attempt to characterise some of the properties of key molecules in this process, we examined both the species specificity and a potential requirement for glycosylation. In the first, we established a positive correlation between species-specificity of pollen tube guidance and their taxonomic distance based on the results acquired through a total of nine wild potato species. This suggests there may be species-specific attractants that can facilitate fertilization by the same or closely-related species. In the second, we observed that a glycosylation enriched fraction of total ovule extracts significantly enhanced pollen tube attraction, indicating this post-translational modification may be important to the process of pollen tube guidance in *S. chacoense*. In an attempt to identify candidate attractants, we performed RNA-sequencing and DGE analyses on different ovule conditions of *S. chacoense*. We identified 284 genes specifically expressed by *S. chacoense* ovules in the presence of an embryo sac during the late stages of embryo sac maturation. Of these, 17 were ScCRPs and classified as candidate attractants. Except for a few which were annotated as *EC1s*, most candidates have unknown function, including 2 genes encoding ECA1 family proteins, 4 genes encoding DEFL family proteins and 6 genes having novel cysteine arrangement unique to *Solanum* species. A *dN/dS* ratio comparison of the 17 ScCRPs and their orthologs in other Solanum species indicated some genes may undergo positive selection and thus potentially act in a species-specific manner. A group of three *ScCRP2s* (*DEFLs*) were cloned, expressed and purified in a bacterial system. Recombinant peptides were tested in functional assays and a moderate level of *in vitro* pollen tube attraction activity was reported. We conclude that secreted proteins are likely involved in ovular pollen tube guidance in *S. chacoense* and that ScCRP2s are prime candidates. For future work, it will be interesting to examine the subcellular localization of *ScCRP2* genes and to test whether fertilization and seed set are impacted in knockout plants. We anticipate the data reported here will contribute to the bigger topic of embryo sac development, pollen-pistil interactions and plant reproduction.

## Supporting information

Supplementary Material-JXB

## Abbreviations

aa: amino acids
A: anthesis
2DBA: two days before anthesis
CRP: cysteine-rich peptides
DEFL: defensin-like peptides
DGE: differential gene expression
ECA1: early culture abundant 1
FRK1: FERTILIZATION-RELATED KINASE 1
SIV: semi-*in vivo*
ScOT: Solanum <u>c</u>hacoense <u>O</u>vule <u>T</u>ranscriptome

## Acknowledgements

We thank Dr. Simon Joly from our institute for his advice regarding the usage of PAML package. We thank Dr. Mario Cappadocia for kindly providing the *Solanum chacoense* V26 genotype. We are most grateful to Dr. Sonia Dorion for her expertise and advice regarding protein extraction as well as protein purification, and to Nicolas Boivin for his technical support.

## Author contributions

DPM, DM: conceptualization; VJ: data curation; YL, VJ: formal analysis; YL, MS: investigation; YL, MS: methodology; DPM, DM: project administration; YL, VJ: resources; VJ: software; DPM, DM: supervision; YL, DSM: validation; YL: visualization; YL: writing – original draft; DM: writing – review & editing. DPM: funding acquisition.

## Conflict of interest

No conflict of interest declared.

## Funding

This work was supported by the Fonds de Recherche du Québec − Nature et technologies (FRQNT: PR-148467) and by the Natural Sciences and Engineering Research Council of Canada (NSERC: RGPIN-03883).

## Data availability

All raw sequence data used in this study have been submitted to the NCBI sequence read archive (http://www.ncbi.nlm.nih.gov/sra) under BioProject PRJNA299204, with accession numbers SRX1370955, SRX1370956, SRX1370981, SRX1370994, SRX1371065, SRX1371085, SRX1371099, SRX1371104-1106, SRX1371110-1115 and SRX1371203-1205. Assembled sequences from the curated hybrid assembly (>200 bp) of *S. chacoense* have been submitted to the Transcriptome Shotgun Assembly Database in GenBank with the accession numbers GEDG01000001-GEDG01042873. All transcripts discussed are associated with a ScOT (<u>S</u>olanum <u>c</u>hacoense <u>O</u>vule <u>T</u>ranscriptome) number which can be used to directly extract their sequences from NCBI Nucleotide Database.

## Notes

https://www.dropbox.com/scl/fo/b649aopyo7wck0oy3jpkq/AFcLnWNviAncjdTa_Bj9bYQ?rlkey=0ifmohhp81nnitvu0fn4o3nh0&st=0qluzxzk&dl=0

### Competing Interest Statement

The authors have declared no competing interest.

