## Supplementary figures and images for "Differential gene expression analysis identifies a group of defensin like peptides from *Solanum chacoense* ovules with *in vitro* pollen tube attraction activity"

### Supplementary Fig S1.jpeg

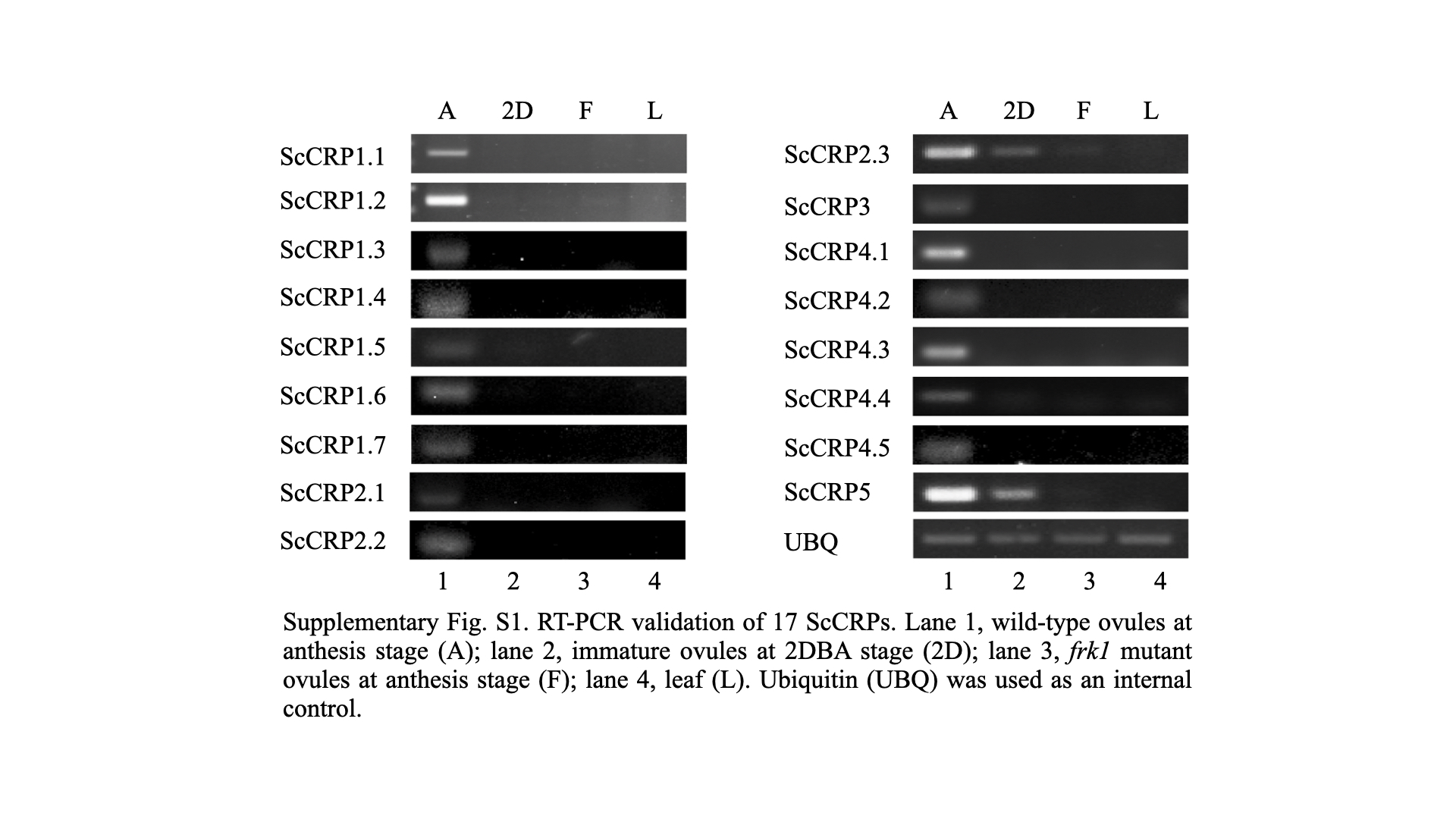

### Supplementary Fig S2.jpeg

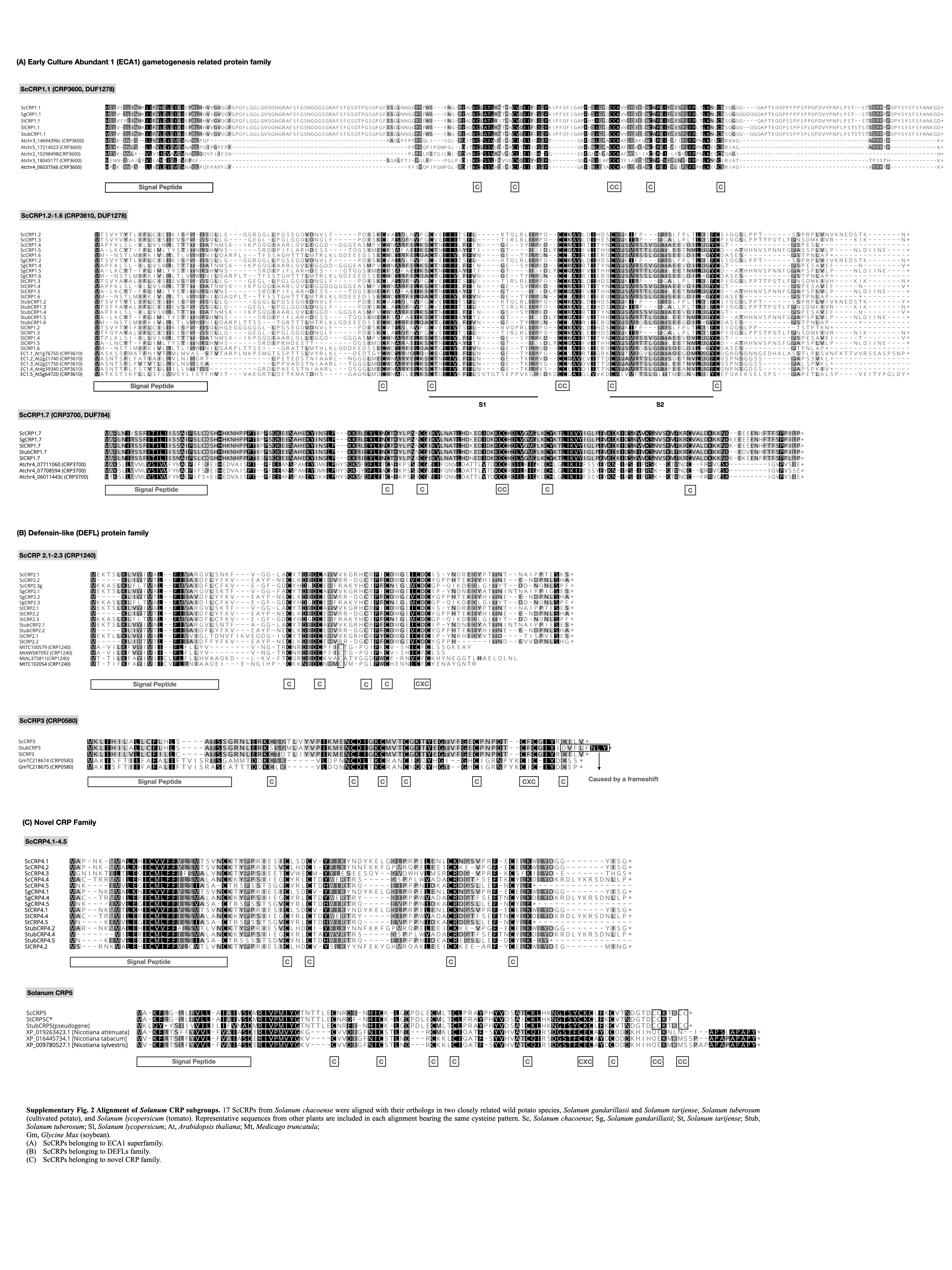
